# Computational mechanisms underlying cortical responses to the affordance properties of visual scenes

**DOI:** 10.1101/177329

**Authors:** Michael F. Bonner, Russell A. Epstein

## Abstract

Biologically inspired deep convolutional neural networks (CNNs), trained for computer vision tasks, have been found to predict cortical responses with remarkable accuracy. However, the complex internal operations of these models remain poorly understood, and the factors that account for their success are unknown. Here we developed a set of techniques for using CNNs to gain insights into the computational mechanisms underlying cortical responses. We focused on responses in the occipital place area (OPA), a scene-selective region of dorsal occipitoparietal cortex. In a previous study, we showed that fMRI activation patterns in the OPA contain information about the navigational affordances of scenes: that is, information about where one can and cannot move within the immediate environment. We hypothesized that this affordance information could be extracted using a set of purely feedforward computations. To test this idea, we examined a deep CNN with a feedforward architecture that had been previously trained for scene classification. We found that the CNN was highly predictive of OPA representations, and, importantly, that it accounted for the portion of OPA variance that reflected the navigational affordances of scenes. The CNN could thus serve as an image-computable candidate model of affordance-related responses in the OPA. We then ran a series of *in silico* experiments on this model to gain insights into its internal computations. These analyses showed that the computation of affordance-related features relied heavily on visual information at high-spatial frequencies and cardinal orientations, both of which have previously been identified as low-level stimulus preferences of scene-selective visual cortex. These computations also exhibited a strong preference for information in the lower visual field, which is consistent with known retinotopic biases in the OPA. Visualizations of feature selectivity within the CNN suggested that affordance-based responses encoded features that define the layout of the spatial environment, such as boundary-defining junctions and large extended surfaces. Together, these results map the sensory functions of the OPA onto a fully quantitative model that provides insights into its visual computations. More broadly, they advance integrative techniques for understanding visual cortex across multiple level of analysis: from the identification of cortical sensory functions to the modeling of their underlying algorithmic implementations.

**AUTHOR SUMMARY:** How does visual cortex compute behaviorally relevant properties of the local environment from sensory inputs? For decades, computational models have been able to explain only the earliest stages of biological vision, but recent advances in the engineering of deep neural networks have yielded a breakthrough in the modeling of high-level visual cortex. However, these models are not explicitly designed for testing neurobiological theories, and, like the brain itself, their complex internal operations remain poorly understood. Here we examined a deep neural network for insights into the cortical representation of the navigational affordances of visual scenes. In doing so, we developed a set of high-throughput techniques and statistical tools that are broadly useful for relating the internal operations of neural networks with the information processes of the brain. Our findings demonstrate that a deep neural network with purely feedforward computations can account for the processing of navigational layout in high-level visual cortex. We next performed a series of experiments and visualization analyses on this neural network, which characterized a set of stimulus input features that may be critical for computing navigationally related cortical representations and identified a set of high-level, complex scene features that may serve as a basis set for the cortical coding of navigational layout. These findings suggest a computational mechanism through which high-level visual cortex might encode the spatial structure of the local navigational environment, and they demonstrate an experimental approach for leveraging the power of deep neural networks to understand the visual computations of the brain.

## INTRODUCTION

Recent advances in the use of deep neural networks for computer vision have yielded image computable models that exhibit human-level performance on scene- and object-classification tasks [1-4]. The units in these networks often exhibit response profiles that are predictive of neural activity in mammalian visual cortex [5-11], suggesting that they might be profitably used to investigate the computational algorithms that underlie biological vision [12-16]. However, many of the internal operations of these models remain mysterious, and the fundamental theoretical principles that account for their predictive accuracy are not well understood [15, 17, 18]. This presents an important challenge to the field: if deep neural networks are to fulfill their potential as a method for investigating visual perception in living organisms, it will first be necessary to develop techniques for using these networks to provide computational insights into neurobiological systems.

It is this issue—the use of deep neural networks for gaining insights into the computational processes of biological vision—that we address here. We focus in particular on the mechanisms underlying natural scene perception. A central aspect of scene perception is the identification of the *navigational affordances* of the local environment—where one can move to (e.g., a doorway or an unobstructed path), and where one’s movement is blocked. In a recent fMRI study, we showed that the navigational-affordance structure of scenes could be decoded from multivoxel response patterns in scene-selective visual areas [19]. The strongest results were found in a dorsal occipital lobe region known as the occipital place area (OPA), which is one of three patches of high-level visual cortex that respond strongly and preferentially to images of spatial scenes [20-24]. These results demonstrated that the OPA encodes affordance-related visual features. However, they did not address the crucial question of how these features might be computed from sensory inputs.

There was one aspect of the previous study that provided a clue as to how affordance representations might be constructed: affordance information was present in the OPA even though participants performed tasks that made no explicit reference to this information. For example, in one experiment, participants were simply asked to report the colors of dots overlaid on the scene, and in another experiment, they were asked to perform a category-recognition task. Despite the fact that these tasks did not require the participants to think about the spatial layout of the scene or plan a route through it, it was possible to decode navigational affordances in the OPA in both cases. This suggested to us that affordances might be rapidly and automatically extracted through a set of purely feedforward computations.

In the current study we tested this idea by examining a biologically inspired CNN with a feedforward architecture that was previously trained for scene classification [3]. This CNN implements a hierarchy of linear-nonlinear operations that give rise to increasingly complex feature representations, and previous work has shown that its internal representations can be used to predict MEG responses to natural scene images [25]. It has also been shown that the higher layers of this CNN can be used to decode the coarse spatial properties of scenes, such as their overall size [25]. By examining this CNN, we aimed to demonstrate that affordance information could be extracted by a feedforward system, and to better understand how this information might be computed.

To preview our results, we find that the CNN contains information about fine-grained spatial features that could be used to map out the navigational pathways within a scene; moreover, these features are highly predictive of affordance-related fMRI responses in the OPA. These findings demonstrate that the CNN can serve as a candidate, image-computable model of navigational-affordance coding in the human visual system. Using this quantitative model, we then develop a set of techniques that provide insights into the computational operations that give rise to affordance-related representations. These analyses reveal a set of stimulus input features that are critical for predicting affordance-related cortical responses, and they suggest a set of high-level, complex features that may serve as a basis set for the population coding of navigational affordances. By combining neuroimaging findings with a fully quantitative computational model, we are able to complement a theory of cortical representation with discoveries of its algorithmic implementation—thus providing insights at multiple levels of understanding and moving us toward a more comprehensive functional description of visual cortex.

## RESULTS

### Representation of navigational affordances in scene-selective visual cortex

To test for the representation of navigational affordances in the human visual system, we examined fMRI responses to 50 images of indoor environments with clear navigational paths passing through the bottom of the scene (Fig. 1A). Subjects viewed these images one at a time for 1.5 s each while maintaining central fixation and performing a category-recognition task that was unrelated to navigation (i.e., press a button when the viewed scenes was a bathroom). Details of the experimental paradigm and a complete analysis of the fMRI responses can be found in a previous report [19]. In this section, we briefly recapitulate the aspects of the results that are most relevant to the subsequent computational analyses.

**Figure 1.**
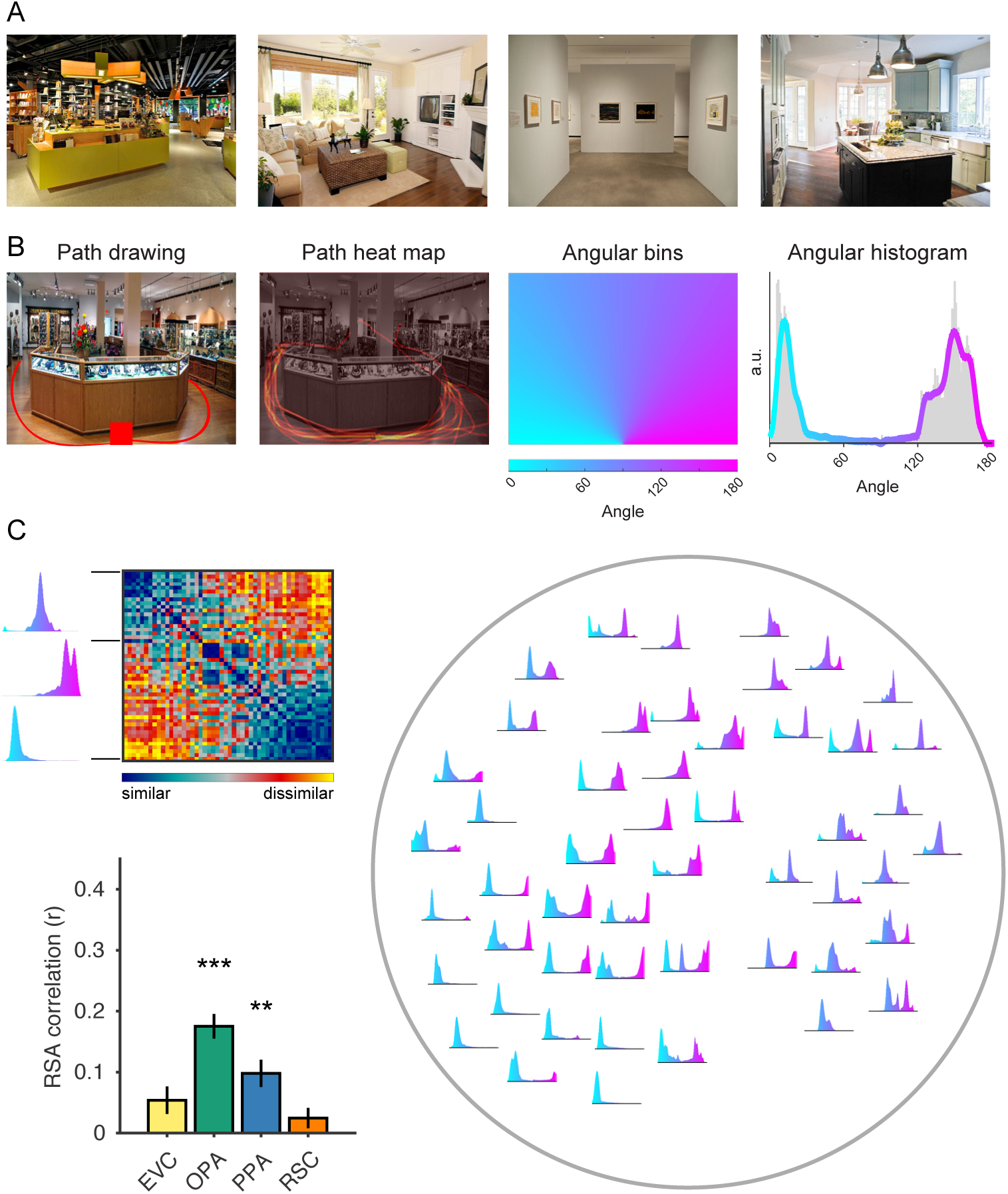
Navigational-affordance information is coded in scene-selective visual cortex. (A) Examples of natural images used in the fMRI experiment. All experimental stimuli were images of indoor environments with clear navigational paths proceeding from the bottom center of the image. (B) In a norming study, an independent group of raters indicated with a computer mouse the paths that they would take to walk through each scene, starting from the red square at the bottom center of the image (far left panel). These data were combined across subjects to create heat maps of the navigational paths in each image (middle left panel). We summed the values in these maps along one-degree angular bins radiating from the bottom center of the image (middle right panel), which produced histograms of navigational probability measurements over a range of angular directions (far right panel). The gray bars in this histogram represent raw data, and the overlaid line indicates the angular data after smoothing. (C) The navigational histograms were compared pairwise across all images to create a model RDM of navigational-affordance coding (top left panel). Right panel shows a two-dimensional visualization of this representational model, created using t-distributed stochastic neighbor embedding (t-SNE), in which the navigational histograms for each condition are plotted within the two-dimensional embedding. RSA correlations were calculated between the model RDM and neural RDMs for each ROI (bottom left panel). The strongest RSA effect for the coding of navigational affordances was in the OPA. There was also a significant effect in the PPA. Error bars represent bootstrap ±1 s.e.m. a.u. = arbitrary units. **p< 0.01, ***p< 0.001

To measure the navigational affordances of these stimuli, we asked an independent group of subjects to indicate with a computer mouse the paths that they would take to walk through each environment starting from the bottom of the image (Fig. 1B). From these responses, we created probabilistic maps of the navigational paths through each scene. We then constructed histograms of these navigational probability measurements in one-degree angular bins over a range of directions radiating from the starting point of the paths. These histograms approximate a probabilistic affordance map of potential navigational paths radiating from the perspective of the viewer [26].

We then tested for the presence of affordance-related information in fMRI responses using representational similarity analysis (RSA) [27]. In RSA, the information encoded in brain responses is compared with a cognitive or computational model through correlations of their representational dissimilarity matrices (RDMs). RDMs are constructed through pairwise comparisons of the model representations or brain responses for all stimulus classes (in this case, the 50 images), and they serve as a summary measurement of the stimulus-class distinctions. The correlation between any two RDMs reflects the degree to which they contain similar information about the stimuli. We constructed an RDM for the navigational-affordance model through pairwise comparisons of the affordance histograms (Fig. 1C). Neural RDMs were constructed for several regions of interest (ROIs) through pairwise comparisons of their multivoxel activation patterns for each image.

We focused our initial analyses on three ROIs that are known to be strongly involved in scene processing: the OPA, the parahippocampal place area (PPA), and the retrosplenial complex (RSC) [20-24]. All three of these regions respond more strongly to spatial scenes (e.g., images of landscapes, city streets, or rooms) than other visual stimuli, such as objects and faces, and thus are good candidates for supporting representations of navigational affordances. We also examined patterns in early visual cortex (EVC). Using RSA to compare the RDMs for these regions to the navigational-affordance RDM, we found evidence that affordance information is encoded in scene-selective visual cortex, most strongly in the dorsal scene-selective region known as the OPA (Fig. 1C). These effects were not observed in lower-level EVC, suggesting that navigational affordances likely reflect mid-to-high-level visual features that require several computational stages along the cortical hierarchy. In our previous report, a whole-brain searchlight analysis confirmed that the strongest cortical locus of affordance coding overlapped with the OPA [19]. Interestingly, affordance coding in scene regions was observed even though participants performed a perceptual-semantic recognition task that did not require them to analyze the navigational affordances of the scene—suggesting that affordance information is automatically elicited during scene perception. Together, these results suggest that scene-selective visual cortex routinely encodes complex spatial features that can be used to map out the navigational affordances of the local visual scene.

These analyses provide functional insights into visual cortex at the level of representation—that is, the identification of sensory information encoded in cortical responses. However, an equally important question for any theory of sensory cortical function is to understand *how* its representations can be computed at an algorithmic level [12-16]. Understanding the algorithms that give rise to high-level sensory representations requires a quantitative model that implements representational transformations from visual stimuli. Thus, we next turn to the question of how affordance representations might be computed from sensory inputs.

### Explaining affordance-related cortical representations with a feedforward image-computable model

Visual cortex implements a complex set of highly nonlinear transformations that remain poorly understood. Attempts at modeling these transformations using hand-engineered algorithms have long fallen short of accurately predicting mid-to-high-level sensory representations [6, 10, 11, 28-30]. However, advances in the development of artificial deep neural networks have dramatically changed the outlook for the quantitative modeling of visual cortex. In particular, recently developed deep CNNs for tasks such as image classification have been found to predict sensory responses throughout much of visual cortex at an unprecedented level of accuracy [5-11]. The performance of these CNNs suggests that they hold the promise of providing fundamental insights into the computational algorithms of biological vision. However, because their internal representations were not hand-engineered to test specific theoretical operations, they are challenging to interpret. Indeed, most of the critical parameters in CNNs are set through supervised learning for the purpose of achieving accurate performance on computer vision tasks, meaning that the resulting features are unconstrained by *a priori* theoretical principles. Furthermore, the complex transformations of these internal CNN units cannot be understood through a simple inspection of their learned parameters. Thus, neural network models have the potential to be highly informative to sensory neuroscience, but a critical challenge for moving forward is the development of techniques to probe the factors that best account for similarities between cortical responses and the internal representations of the models.

Here we tested a deep CNN as a potential candidate model of affordance-related responses in scene-selective visual cortex. Given the apparent automaticity of affordance-related responses, we hypothesized that they could be modeled through a set of purely feedforward computations performed on image inputs. To test this idea, we examined a model that was previously trained to classify images into a set of scene categories [3]. This feedforward model contains 5 convolutional layers followed by 3 fully connected layers, the last of which contains units corresponding to a set of scene category labels (Fig. 2A). The architecture of the model is highly similar to the AlexNet model that initiated the recent surge of interest in CNNs for computer vision [2]. Units in the convolutional layers of this model have local connectivity, giving rise to increasingly large spatial receptive fields from layers 1 through 5. The dense connectivity of the final three layers means that the selectivity of their units could depend on any spatial position in the image. Each unit in the CNN implements a linear-nonlinear operation in which it computes a weighted linear sum of its inputs followed by a nonlinear activation function (specifically, a rectified linear threshold). The weights on the inputs for each unit define a type of filter, and each convolutional layer contains a set of filters that are replicated with the same set of weights over all parts of the image (hence, the term “convolution”). There are two other nonlinear operations implemented by a subset of the convolutional layers: max-pooling, in which only the maximum activation in a local pool of units is passed to the next layer, and normalization, in which activations are adjusted through division by a factor that reflects the summed activity of multiple units at the same spatial position. Together, this small set of functional operations along with a set of architectural constraints define an untrained model whose many other parameters can be set through gradient descent with backpropagation—producing a trained model that performs highly complex feats of visual classification.

**Figure 2.**
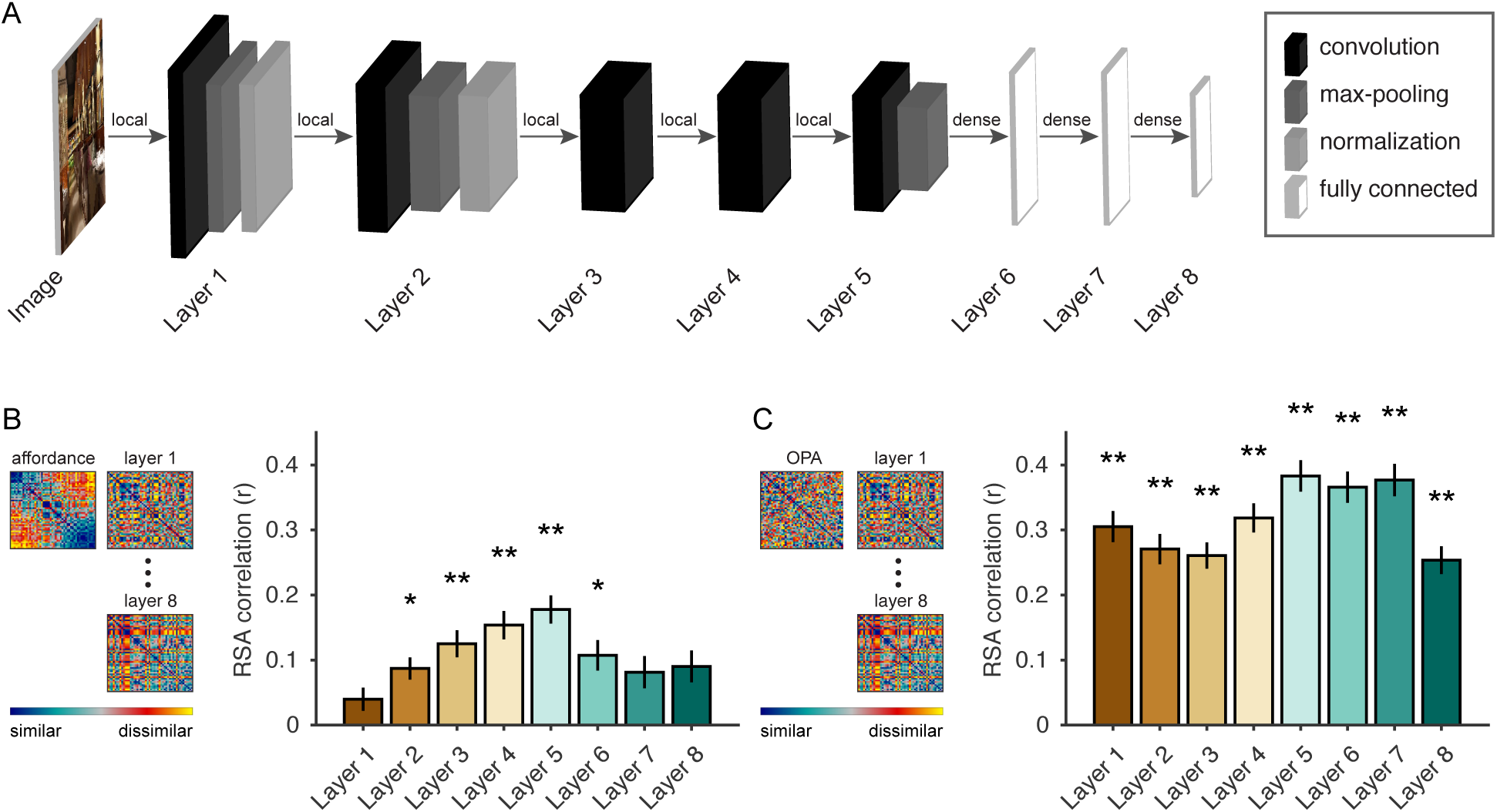
Navigational-affordance information can be extracted by a feedforward computational model. (A) Architecture of a deep CNN trained for scene categorization. Image pixel values are passed to a feedforward network that performs a series of linear-nonlinear operations, including convolution, rectified linear activation, local max pooling, and local normalization. The final layer contains category-detector units that can be interpreted as signaling the association of the image with a set of semantic labels. (B) RSA of the navigational-affordance model and the outputs from each layer of the CNN. The affordance model correlated with multiple layers of the CNN, with the strongest effects observed in higher convolutional layers and weak or no effects observed in the earliest layers. This is consistent with the findings of the fMRI experiment, which indicate that navigational affordances are coded in mid- to-high-level visual regions but not early visual cortex. (C) RSA of responses in the OPA and the outputs from each layer of the CNN. All layers showed strong RSA correlations with the OPA, and the peak correlation was in layer 5, the highest convolutional layer. Error bars represent bootstrap ±1 s.e.m. *p< 0.05, **p< 0.01

We passed the images from the fMRI experiment through the CNN and constructed a set of RDMs using the final outputs from each layer. We then used RSA to compare the representations of the CNN with: (i) the RDM for the navigational-affordance model and (ii) the RDM for fMRI responses in the OPA. The RSA comparisons with the affordance model showed that the CNN contained affordance-related information, which arose gradually across the lower layers and peaked in layer 5, the highest convolutional layer (Fig. 2B). The weak effects in lower convolutional layers are consistent with the pattern of findings from the fMRI experiment, in which affordance representations were not evident in EVC, and they suggest that affordances reflect mid-to-high-level, rather than low-level, visual features. The decrease in affordance-related information in the last three fully connected layers may result from the increasingly semantic nature of representations in these layers, which ultimately encode a set of scene-category labels that are likely orthogonal to the affordance-related features of the scenes. The RSA comparisons with OPA responses showed that the CNN provided a highly accurate model of representations in this brain region, with strong effects across all CNN layers and a peak correlation in layer 5 (Fig. 2B). Indeed, several layers of the CNN reached the highest accuracy we could expect for any model, given the noise ceiling of the OPA, which was calculated from the variance across subjects (r-value for OPA noise ceiling = 0.30). Together, these findings demonstrate the feasibility of computing complex affordance-related features through a set of purely feedforward transformations, and they show that the CNN is a highly predictive model of OPA responses to natural images depicting such affordances.

The above findings demonstrate that the CNN is representationally similar to the navigational-affordance RDM and also similar to the OPA RDM, but they leave open the important question of whether the CNN captures the same variance in the OPA as the navigational-affordance RDM. In other words, can the CNN serve as a computational model for *affordance-related* responses in the OPA? To address this question, we combined the RSA approach with commonality analysis [31], a variance partitioning technique in which the explained variance of a multiple regression model is divided into the unique and shared variance contributed by all its predictors. In this case, multiple regression RSA was used to construct an encoding model of OPA representations. Thus, the OPA was the predictand and the affordance and CNN models were predictors. Our goal was to identify the portion of the shared variance between the affordance RDM and OPA RDM that could be accounted for by the CNN RDM (Fig. 3A). This analysis showed that the CNN could explain a substantial portion of the representational similarity between the navigational-affordance model and the OPA. In particular, over half of the explained variance of the navigational-affordance RDM could be accounted for by layer 5 of the CNN (Fig. 3B). This suggests that the CNN can serve as a candidate, quantitative model of affordance-related responses in the OPA.

**Figure 3.**
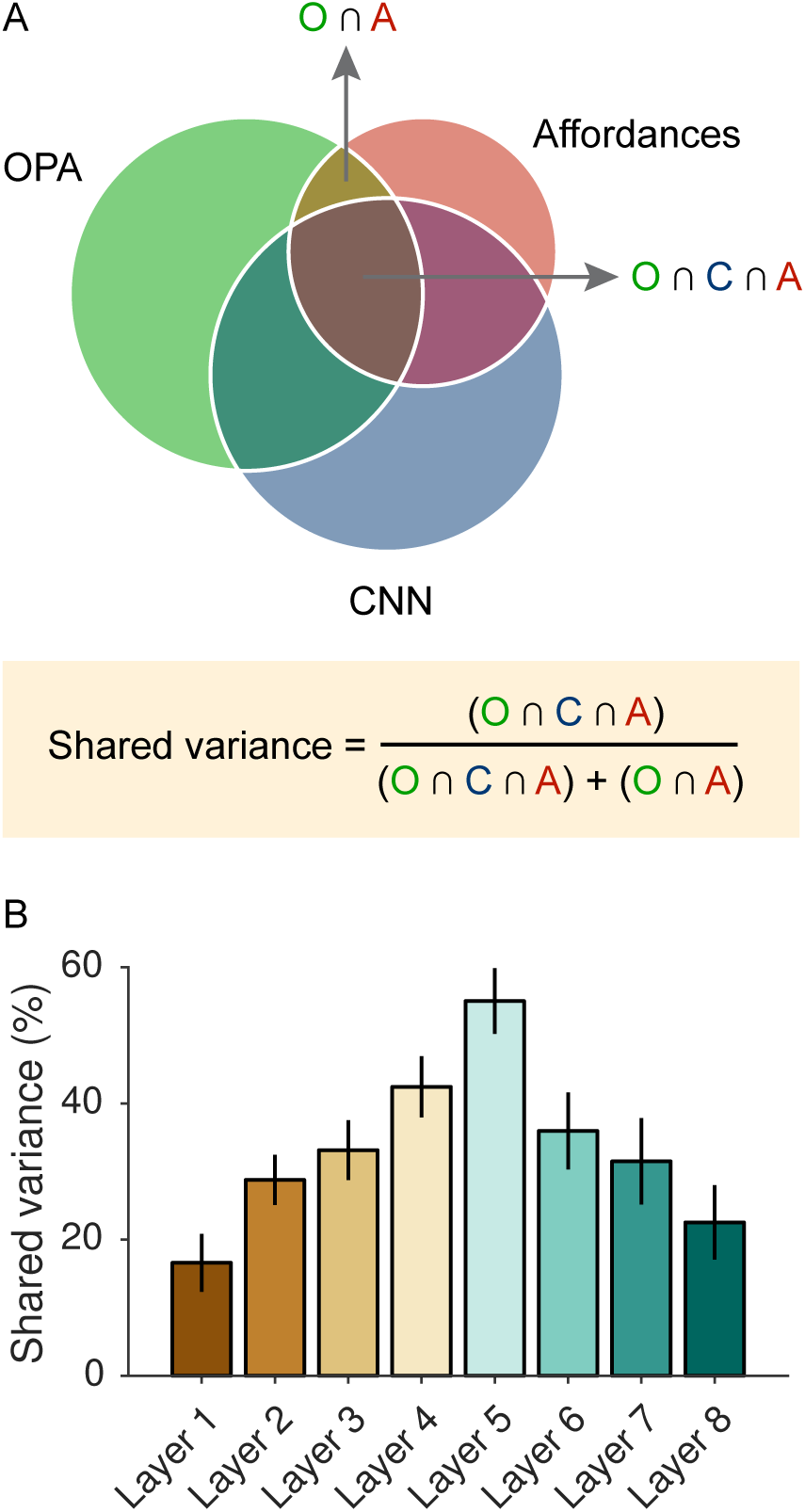
The CNN accounts for shared variance between OPA responses and the navigational-affordance model. (A) A variance-partitioning procedure, known as commonality analysis, was used to quantify the portion of the shared variance between the OPA RDM and the navigational-affordance RDM that could be accounted for by the CNN. Commonality analysis partitions the explained variance of a multiple regression model into the unique and shared variance contributed by all its predictors. In this case, multiple regression RSA was performed with the OPA as the predictand and the affordance and CNN models as predictors. (B) Partitioning the explained variance of the affordance and CNN models showed that over half of the variance explained by the navigational-affordance model in the OPA could be accounted for by the highest convolutional layer of the CNN (layer 5). Error bars represent bootstrap ±1 s.e.m.

One of the most important aspects of the CNN as a candidate model of affordance-related cortical responses is that it is image computable, meaning that its representations can be calculated for any input image. This makes it possible to test predictions about the internal computations of the model by generating new stimuli and running *in silico* experiments. In the next two sections, we run a series of experiments on the CNN to gain insights into the factors that underlie its predictive accuracy in explaining the representations of the navigational-affordance model and the OPA.

### Low-level image features that underlie the predictive accuracy of the computational model

A fundamental issue for understanding any model of sensory computation is determining the aspects of the sensory stimulus on which it operates. In other words, what sensory inputs drive the responses of the model? Here we investigated the image features that drive affordance-related responses in the CNN. Specifically, we sought to identify classes of low-level stimulus features that are critical for explaining the representational similarity of the CNN to the navigational-affordance model and the OPA.

We expected that navigational affordances would rely on image features that convey information about the spatial structure of scenes. Our specific hypotheses were that affordance-related representations would be relatively unaffected by color information and would rely heavily on high spatial frequencies and edges at cardinal orientations (i.e., horizontal and vertical). The hypothesis that color information would be unimportant was motivated by our intuition that color is not typically a defining feature of the structural properties of scenes and by a previous finding of ours showing that affordance representations in the OPA are partially tolerant to variations in scene textures and colors [19]. The other two hypotheses were motivated by previous work suggesting that high spatial frequencies and cardinal orientations are especially informative for the perceptual analysis of spatial scenes, and that the PPA and possibly other scene-selective regions are particularly sensitive to these low-level visual features [32-36], but see [37].

To test these hypotheses, we generated new sets of filtered stimuli in which specific visual features were isolated or removed (i.e., color, spatial frequencies, cardinal or oblique edges; Fig. 4A-B). These filtered stimuli were passed through the CNN, and new RDMs were created for each layer. We used the commonality-analysis technique described in the previous section to quantify the portion of the original explained variance of the CNN that could be accounted for by the filtered stimuli. This procedure was applied to the explained variance of the CNN for predicting both the navigational-affordance RDM and the OPA RDM (Fig. 4A). The results for both sets of analyses showed that over half of the explained variance of the CNN could be accounted for when the inputs contained only grayscale information, high-spatial frequencies, or edges at cardinal orientations. In contrast, when input images with low-spatial frequencies and oblique edges were used, a much smaller portion of the explained variance was accounted for. The differences in explained variance across high and low spatial frequencies and across cardinal and oblique orientations were more pronounced for the RSA predictions of the affordance RDM, but a similar pattern was observed for the OPA RDM. We used a bootstrap resampling procedure to estimate 95% confidence intervals for these sets of comparisons. The confidence bounds from this analysis showed that the differences in shared variance for high vs. low spatial frequencies and for cardinal vs. oblique orientations were reliable for both the affordance RDM and the OPA RDM.

**Figure 4.**
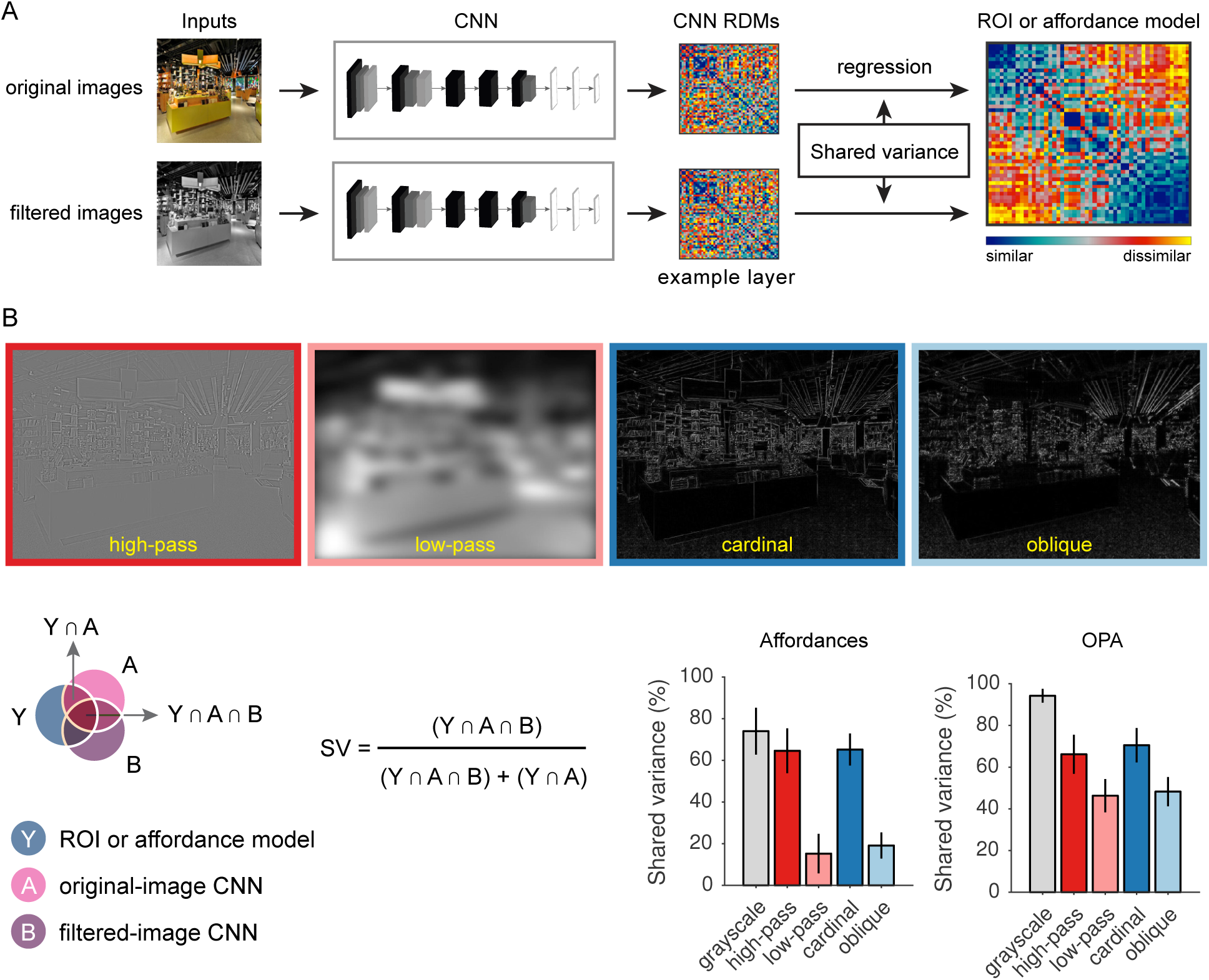
Analysis of low-level image features that underlie the predictive accuracy of the CNN. (A) Experiments were run on the CNN to quantify the contribution of specific low-level image features to the representational similarity between the CNN and the OPA and between the CNN and the navigational-affordance model. First, the original stimuli were passed through the CNN, and RDMs were created for each layer. Then the stimuli were filtered to isolate or remove specific visual features. For example, grayscale images were created to remove color information. These filtered stimuli were passed through the CNN, and new RDMs were created for each layer. Multiple-regression RSA was performed using the RDMs for the original and filtered stimuli as predictors. Commonality analysis was applied to this regression model to quantify the portion of the shared variance between the CNN RDM and the OPA RDM or between the CNN RDM and the affordance RDM that could be accounted for by the filtered stimuli. (B) This procedure was used to quantify the contribution of color (grayscale), spatial frequencies (high-pass and low-pass), and edge orientations (cardinal and oblique). The RSA effects of the CNN were driven most strongly by grayscale information at high spatial frequencies and cardinal orientations. Over half of the shared variance between the CNN and the OPA and between the CNN and the affordance model could be accounted for by representations of grayscale images or images containing only high-spatial frequency information or edges at cardinal orientations. In contrast, the contribution of low spatial frequencies and edges at oblique orientations were considerably lower. These differences in high-versus-low spatial frequencies and cardinal-versus-oblique orientations were more pronounced for RSA predictions of the navigational-affordance RDM, but a similar pattern was observed for the OPA RDM as well. Bars represent means and error bars represent ±1 s.e.m. across CNN layers.

Together, these results suggest that visual inputs at high-spatial frequencies and cardinal orientations are critical for computing the affordance-related features of the CNN. Furthermore, these computational operations appear to be largely tolerant to the removal of color information. Indeed, it is striking how much explained variance these inputs account for given how much information has been discarded from their corresponding filtered stimulus sets.

### Visual-field biases that underlie the predictive accuracy of the computational model

In addition to examining classes of input features to the CNN, we also sought to understand how inputs from different spatial positions in the image affected the similarity between the CNN and RDMs for the navigational-affordance model and the OPA. Our hypothesis was that these RSA effects would be driven most strongly by inputs from the lower visual field (we use the term “visual field” here because the fMRI subjects were asked to maintain central fixation throughout the experiment). This was motivated by previous findings showing that the OPA has a retinotopic bias for the lower visual field [38, 39] and the intuitive prediction that the navigational affordances of local space rely heavily on features close to the ground plane.

To test this hypothesis, we generated sets of occluded stimuli in which everything except a small horizontal slice of the image was masked (Fig. 5). These occluded stimuli were passed through the CNN, and new RDMs were created for each layer. Once again, we used the commonality-analysis technique described above to quantify the portion of the original explained variance of the CNN that could still be accounted for by these occluded stimuli. This procedure was repeated with the un-occluded region slightly shifted on each iteration until the entire vertical axis of the image was sampled. We used this procedure to analyze the explained variance of the CNN for predicting both the navigational-affordance RDM and the OPA RDM (Fig. 5). For comparison, we also applied this procedure to RDMs for the other ROIs. These analyses showed that the predictive accuracy of the CNN for both the affordance model and the OPA were driven most strongly by inputs from the lower visual field. Strikingly, as much as 70% of the explained variance of the CNN in the OPA could be accounted for by a small horizontal band of features at the bottom of the image (Fig. 5). We created a summary statistic for this visual-field bias by calculating the difference in mean shared variance across the lower and upper halves of the image. A comparison of this summary statistic across all tested RDMs shows that the lower visual field bias was observed only for RSA predictions of the affordance model and the OPA, but not the other ROIs (Fig. 5).

**Figure 5.**
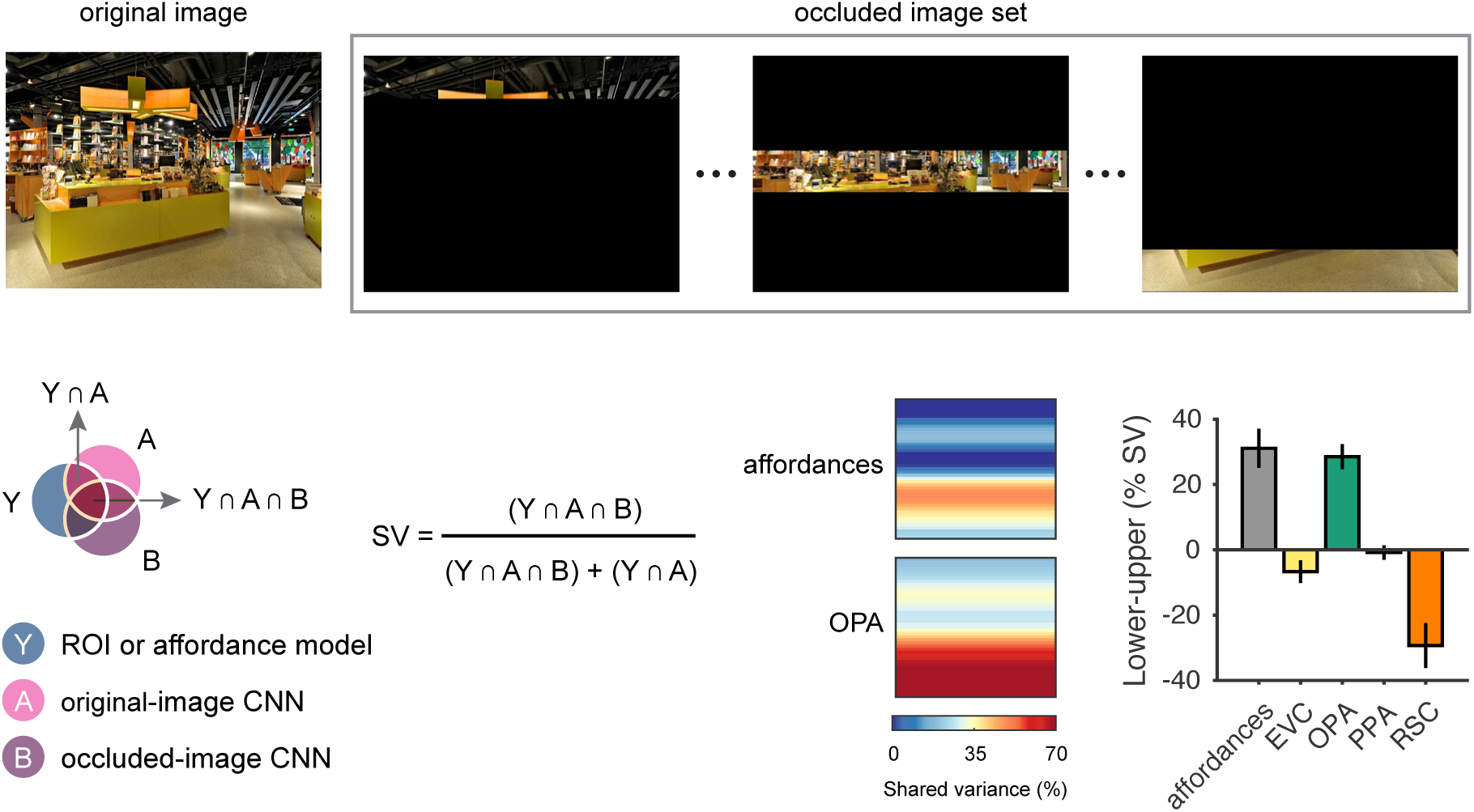
Visual-field biases in the predictive accuracy of the CNN. Experiments were run on the CNN to quantify the importance of visual inputs at different positions along the vertical axis of the image. First, the original stimuli were passed through the CNN, and RDMs were created. Then the stimuli were occluded to mask everything outside of a small horizontal slice of the image (top panel). These occluded stimuli were passed through the CNN, and new RDMs were created. Multiple regression RSA was performed using the RDMs for the original and occluded images as predictors. Commonality analysis was applied to this regression model to quantify the portion of the shared variance between the CNN and the OPA or between the CNN and the navigational-affordance model that could be accounted for by the occluded images (bottom left panel). This procedure was repeated with the un-occluded region slightly shifted on each iteration until the entire vertical axis of the image was sampled. Results indicated that RSA effects of the CNN were driven most strongly by features in the lower half of the image (bottom right panel). This effect was most pronounced for RSA predictions of the OPA RDM, in which ∼70% of the explained variance of the CNN could be accounted for by visual information within a small slice of the image from the lower visual field. A summary statistic of this visual-field bias, created by calculating the difference in mean shared variance across the lower and upper halves of the image, showed that a bias for information in the lower visual field was observed for the affordance model and the OPA, but not for EVC, PPA, or RSC. Bars represent means and error bars represent ±1 s.e.m. across CNN layers.

Together, these results demonstrate that information from the lower visual field is critical to the performance of the CNN in predicting the affordance RDM and the OPA RDM. These findings are consistent with previous neuroimaging work on the retinotopic biases of the OPA [38, 39], and they suggest that the cortical computation of affordance-related features reflects a strong bias for inputs from the lower visual field.

### High-level features of the computational model that best account for affordance-related cortical representations

The analyses above examined the stimulus inputs that drive affordance-related computations in the CNN. We next set out to characterize the high-level features that result from these computations. Specifically, we sought to characterize the internal representations of the CNN that best account for the representations of the OPA and the navigational-affordance model. To do this, we performed a set of visualization analyses that help reify the complex visual motifs detected by the internal units of the CNN.

We characterized the feature selectivity of CNN units using a receptive-field mapping procedure (Fig. 6A) [40]. The goal was to identify natural image features that drive the internal representations of the CNN. In this procedure, the selectivity of individual CNN units was mapped across each image by iteratively occluding the inputs to the CNN. First, the original, un-occluded image was passed through the CNN. Then a small portion of the image was occluded with a patch of random pixel values (11 pixels by 11 pixels). The occluded image was passed though the CNN, and the discrepancies in unit activations relative to the original image were logged. After iteratively applying this procedure across all spatial positions in the image, a two-dimensional discrepancy map was generated for each unit and each image (Fig. 6A). Each discrepancy map indicates the sensitivity of a CNN unit to the visual information across all spatial positions of an image. The spatial distribution of the discrepancy effects reflects the position and extent of a unit’s receptive field, and the magnitude of the discrepancy effects reflects the sensitivity of a unit to the underlying image features. We focused our analyses on the units in layer 5, which was the layer with the highest RSA correlation for the both the navigational-affordance model and the OPA. We selected 50 units in this layer based on their unit-wise RSA correlations to the navigational-affordance model and the OPA. These units were highly informative for our effects of interest: an RDM created from just these 50 units showed comparable RSA correlations to those observed when using all units in layer 5 (correlation with affordance RDM: r = 0.28; correlation with OPA RDM: r = 0.35). We generated receptive-field visualizations for each of these units. These visualizations were created by identifying the top 3 images that drove the responses within a unit’s receptive field. A segmentation mask was then applied to each image by thresholding the unit’s discrepancy map at 10% of the peak discrepancy value. Segmentations highlight the portion of the image that the unit was sensitive to. Each segmentation is outlined in red, and regions of the image outside of the segmentation are darkened (Fig. 6B).

**Figure 6.**
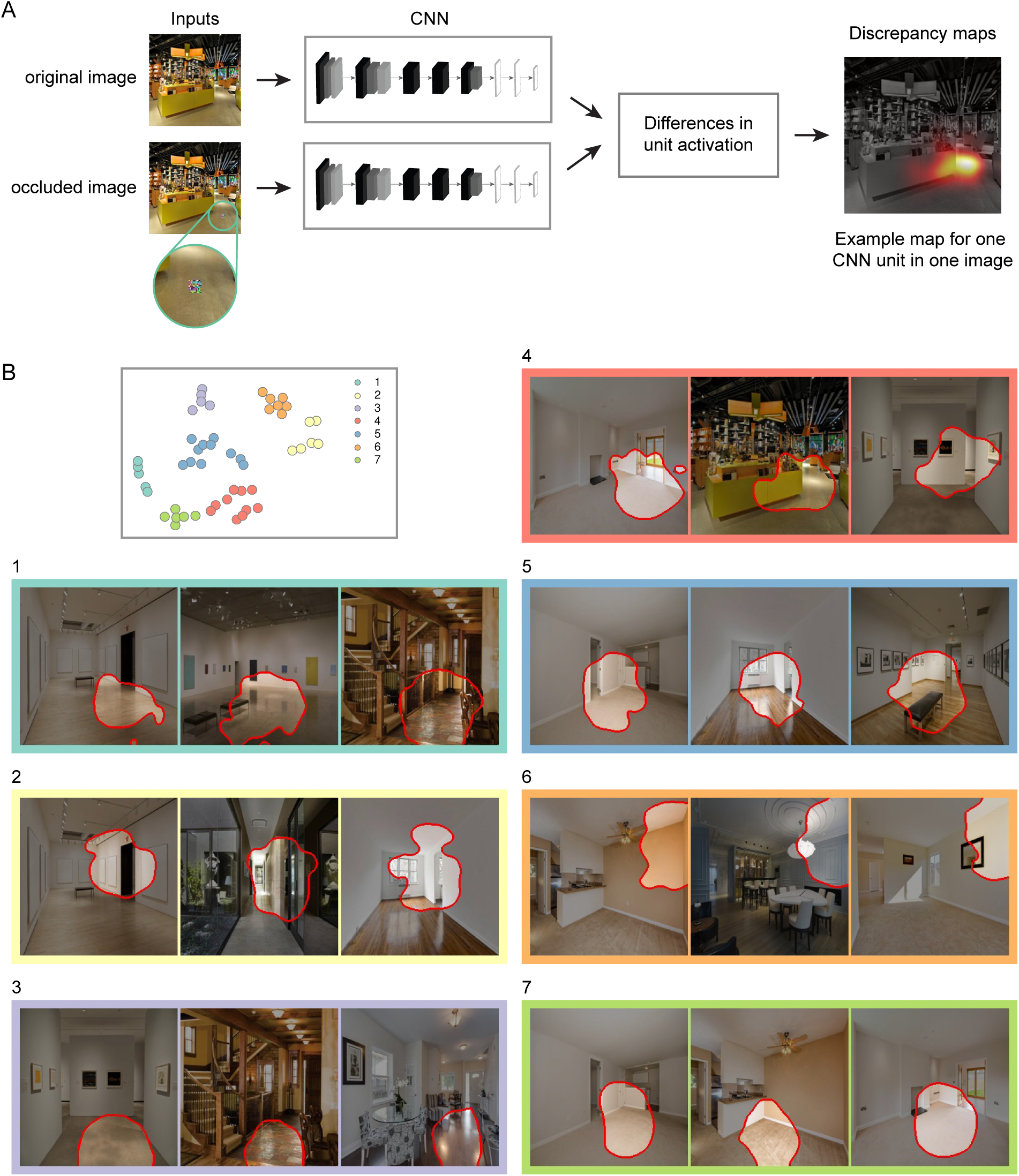
Receptive-field selectivity of CNN units. (A) The selectivity of individual CNN units was mapped across each image through an iterative occlusion procedure. First, the original image was passed through the CNN. Then a small portion of the image was occluded with a patch of random pixel values. The occluded image was passed though the CNN, and the discrepancies in unit activations relative to the original image were logged. After iteratively applying this procedure across all spatial positions in the image, a two-dimensional discrepancy map was generated for each CNN unit and each stimulus (far right panel). Each discrepancy map indicates the sensitivity of a CNN unit to the visual information within an image. The two-dimensional position of its peak effect reflects the unit’s spatial receptive field, and the magnitude of its peak effect reflects the unit’s selectivity for the image features within this receptive field. (B) Receptive-field visualizations were generated for a subset of the units in layer 5 that had strong unit-wise RSA correlations with the OPA and the affordance model. To examine the visual motifs detected by these units, we created a two-dimensional embedding of the units based on the visual similarity of the image features that drove their responses. A clustering algorithm was then used to identify groups of units whose responses reflect similar visual motifs (top left panel). This data-driven procedure identified 7 clusters, which are color-coded and numbered in the two-dimensional embedding. Visualizations are shown for an example unit from each cluster (the complete set of visualizations can be seen in Fig. S1-S7). These visualizations were created by identifying the top 3 images that drive the responses within a unit’s receptive field. A segmentation mask was then applied to each image by thresholding the unit’s discrepancy map at 10% of the peak discrepancy value. Segmentations highlight the portion of the image that the unit was sensitive to. Each segmentation is outlined in red, and regions of the image outside of the segmentation are darkened. Among these visualizations, two broad themes were discernable: boundary-defining junctions (e.g., clusters 1, 5, 6, and 7) and large extended surfaces (e.g., cluster 3). The boundary-defining junctions included junctions where two or more large planes meet (e.g., a wall and a floor). Large extended surfaces included uninterrupted portions of floor and wall planes. There were also units that detected features indicative of doorways and other open pathways (e.g., clusters 2 and 4). All of these high-level features appear to be well-suited for mapping out the spatial layout and navigational boundaries in a visual scene.

We sought to identify prominent trends across this set of receptive-field segmentations. In a simple visual inspection of the segmentations, we detected visual motifs that were common among the units, and the results of an automated clustering procedure highlighted these trends. Using data-driven techniques, we embedded the segmentations into a low-dimensional projection and then partitioned them into clusters with similar visual motifs. We used t-distributed stochastic neighbor embedding (t-SNE) to generate a two-dimensional projection of the units based on the visual similarity of their receptive-field segmentations (Fig. 6B). We then used k-means clustering to identify sets of units with similar embeddings. The number of clusters was set at 7 based on the outcome of a cluster-evaluation procedure. The specific cluster assignments do not necessarily indicate major qualitative distinctions between units. Rather, they provide a data-driven means of reducing the complexity of the results and highlighting the broad themes in the data. These themes can also be seen in the complete set of visualizations plotted in Fig. S1-S7.

These visualizations revealed two broad visual motifs: boundary-defining junctions and large, extended surfaces. Boundary-defining junctions are the regions of an image where two or more extended planes meet (e.g., clusters 1, 5, 6, and 7 in Fig. 6B). These were often the junctions of walls and floors, and less often ceilings. This was the most common visual motif across all segmentations. Large, extended surfaces were uninterrupted portions of floor and wall planes (e.g., cluster 3 in Fig. 6B). There were also units that detected more complex structural features that were often indicative of doorways and other open pathways (e.g., clusters 2 and 4 in Fig. 6B).

A common thread running through all these visualizations is that they appear to reflect high-level scene features that could be reliably used to map out the spatial layout and navigational affordances of the local environment. Boundary-defining junctions and large, extended surfaces provide critical information about the spatial geometry of the local scene, and more fine-grained structural elements, such as doorways and open pathways, are critical to the navigational layout of a scene. Together, these results suggest a minimal set of high-level visual features that are critical for modeling the navigational affordances of natural images and predicting the affordance-related responses of scene-selective visual cortex.

## DISCUSSION

We examined a deep CNN from computer vision for insights into the computations of high-level visual cortex during natural scene perception. Previous work has shown that supervised CNNs, trained for tasks such as image classification, are predictive of sensory responses throughout much of visual cortex, but their internal operations are highly complex and remain poorly understand. Here we developed a set of techniques for relating the internal operations of a CNN to cortical sensory functions. Our approach combines hypothesis-driven *in silico* experiments with statistical tools for quantifying the shared representational content in neural and computational systems. We applied these techniques to understand the computations that give rise to navigational-affordance representations in the OPA. We found that affordance-related cortical responses could be predicted through a set of purely feedforward computations, involving several stages of nonlinear feature transformations. These computations relied heavily on high-spatial-frequency information at cardinal orientations, and were most strongly driven by inputs from the lower visual field. Following several computational stages, this model gives rise to large, complex features that convey information about the structural layout of a scene. Visualization analyses suggested that the most prominent motifs among these high-level features were the junctions and surfaces of extended planes found on walls, floors, and large objects. Together, these results identify a biologically plausible set of feedforward computations that account for a critical function of high-level visual cortex, and they shed light on the stimulus features that drive these computations and the internal representations that they give rise to.

### Information processing in scene-selective cortex

These findings have important implications for developing a computational understanding of scene-selective visual cortex. To gain an understanding of the algorithms implemented by the visual system we first need candidate quantitative models whose parameters and operations can be interpreted for theoretical insights. One of the primary criteria for evaluating such a model is that it explains a substantial portion of stimulus-driven activity in the brain region of interest. Any model that does not meet this necessary criterion is fundamentally insufficient or incorrect. A major strength of the CNN examined here is that it is highly accurate at predicting cortical responses to the perception of natural scenes. Indeed, the CNN explained as much variance in the responses of the OPA as could be expected for any model, after accounting for the portion of OPA variance that could be attributed to noise. Thus, as in previous studies of high-level object perception, the ability of the CNN to reach the noise ceiling for explained variance during scene perception constitutes a major advance in the quantitative modeling of cortical responses [6, 7].

Another strength of the CNN as a candidate model is that its representations can be computed from arbitrary image inputs. This image computability confers two major benefits to the CNN. First, its internal representations can be investigated across all computational stages and mapped onto a cortical hierarchy, allowing for a complete description of the nonlinear transformations that convert sensory inputs into high-level visual features [11]. Second, image computability allows investigators to submit novel stimulus inputs to the CNN for the purpose of testing mechanistic hypotheses through *in silico* experiments. Here we took advantage of this image computability to test several hypotheses about which stimulus inputs are critical for computing affordance-related visual features and predicting the responses of the OPA. These analyses demonstrated the importance of inputs from the lower visual field (i.e., the bottom of the image when fixation is at the center), which aligns with previous fMRI studies that used receptive-field mapping to identify a lower-field bias in the OPA [38, 39]. These analyses also demonstrated the importance of several low-level image features that have previously been shown to drive the responses of scene-selective visual cortex, including high-spatial frequencies and contours at cardinal orientations [32-36], but see [37].

We also performed visualization experiments on the internal representations of the CNN to identify potential affordance-related scene features that might be encoded in the population responses of the OPA. Our approach involved data-driven visualizations of the image regions detected by individual CNN units. We focused on the fifth convolutional layer of the CNN and, in particular, on units in this layer that corresponded most strongly to the representations of the OPA and the navigational-affordance model. Among the scene features detected by these units, two broad themes were prominent: boundary-defining junctions and large, extended surfaces. Boundary-defining junctions were contours where two or more large and often orthogonal surfaces were adjoined. These included extended junctions of two surfaces, such as a wall and a floor, and corners where three surfaces come together, such as two walls and a floor. In other words, boundary-defining junctions resembled the basic features that one would use to sketch the spatial layout of a scene.

The idea that such features might be encoded in scene-selective visual cortex accords with previous findings from neuroimaging and electrophysiology. The most directly related findings come from a series of neuroimaging studies investigating the responses of scene-selective cortex to line drawings of natural images [41, 42]. These line-drawing stimuli convey information about contours and their spatial arrangement, but they lack many of the rich details of natural images, including color, texture, and shading. Nonetheless, these stimuli elicit representations of scene-category information in scene-selective visual cortex. These effects appear to be driven mostly by long contours and their junctions, whose arrangement conveys information about the spatial structure of a scene. This suggests that a substantial portion of the features encoded by scene-selective cortex can be computed using only structure-defining contour information. This aligns with the findings from our visualization analyses, which suggest that large surface junctions are an important component of the information encoded by scene-selective visual cortex. These surface junctions correspond to the long structure-defining contours that would be highlighted in a line drawing. Electrophysiological investigations of scene-selective visual cortex in the macaque brain have also demonstrated the importance of structure-defining contours [43], and even identified cells that exhibited selectivity for the surface junctions in rooms, which appears to be remarkably similar to the selectivity for boundary-defining junctions identified here. Our findings are also broadly consistent with previous behavioral studies demonstrating that contours and contour junctions in 2D images are highly informative about the arrangement of surfaces in 3D space [44]. In particular, contour junctions convey information about 3D structure that is largely invariant to changes in viewpoint, making them exceptionally useful for inferring spatial structure [44].

In addition to boundary-defining junctions, we also observed selectivity for large extended surfaces. One recent study has suggested that the responses of scene-selective visual cortex can be well predicted from the depth and orientation of large surfaces in natural scenes [45]. This appears to be consistent with our finding, but other possible interpretations include texture identification or the use of texture gradients as 3D orientation cues [46, 47]. Together, this pattern of selectivity for large surfaces, in combination with selectivity for the junctions between these surfaces, is consistent with an information-processing mechanism for representing the spatial structure of the local visual environment [12].

Taken as a whole, these computational findings have implications for interpreting previous neuroimaging studies of scene-selective cortex. It has been argued that the apparent category selectivity of scene regions can be explained more parsimoniously in terms of preferences for low-level image features, such as high spatial frequencies [32-36], but see [37]. However, the analyses presented here suggest an alternative interpretation, namely that scene-selective visual regions encode complex features that convey information about high-level scene properties, such as navigational layout, but that the computations that give rise to these features rely heavily on specific sets of low-level inputs [48]. This account characterizes the function of scene-selective visual cortex within the context of a computational system, and it demonstrates how a region within this system could exhibit response preferences for the low-level features that drive its upstream inputs. Thus, by examining a candidate computational model, we identified a potential mechanism through which neuroimaging studies could produce seemingly contradictory findings on the feature selectivity of scene-selective cortex. More broadly, these analyses demonstrate the importance of building explicit computational models to evaluate functional theories of high-level visual cortex. Doing so allows investigators to interpret cortical processes in terms of their functional significance to systems-level computations rather than region-specific representational models.

### Using deep CNNs to obtain insights into biological vision

The analyses and techniques presented here are broadly relevant to research on the functions of visual cortex. A major goal of visual neuroscience is to understand the information-processing mechanisms that visual cortex carries out [12]. Progress toward this goal can be assessed by how well investigators are able to implement these mechanisms *de novo* using models that reflect a compact set of theoretical principles. To this end, investigators require models that are constructed from mathematical algorithms, to allow for implementations in any suitable computational hardware, and whose internal operations are theoretically interpretable, in the sense that one can provide summary descriptions of the functions they carry out and the theoretical principles they embody. A long line of work in visual neuroscience has attempted to understand the information-processing mechanisms of visual cortex by hand-engineering computational models based on *a priori* theoretical principles [28, 29, 49]. Although this approach has been fruitful in characterizing the earliest stages of visual processing, it has not proved effective for explaining the functions of mid-to-high-level visual cortex, where the complexity of the operations and the number of possible features grows exponentially [11]. Recent advances in the development of deep CNNs trained for computer vision have incidentally yielded quantitative models that are remarkably accurate at predicting functional activity throughout much of the visual system [5-11]. However, from a theoretical perspective, these highly complex models have remained largely opaque, and little is known about what aspects of these models might be relevant for understanding the information processes of biological vision.

Here we developed an approach for probing the internal operations of a CNN for insights into cortical computation. Our approach uses RSA in the context of multiple linear regression to evaluate similarities between computational, theoretical, and neural systems. A major benefit of RSA is that it evaluates the information in these systems through summary representations in RDMs, which avoids the many difficulties of identifying mappings between the individual units of high-dimensional systems [27]. We evaluated these multiple linear regressions using a variance-partitioning procedure that allowed us to quantify the degree to which representational models explained shared or unique components of the information content in a cortical region. These statistical methods were combined with techniques for running *in silico* experiments to test theoretically motivated hypotheses about information processing in the CNN and its relationship to the functions of visual cortex. Although we applied these techniques to an exploration of affordance coding in visual scenes, they are broadly applicable and could be used to examine any cortical region or image-computable model. They demonstrate a general approach for exploring how the computations of a CNN relate to the information-processing algorithms of biological vision.

It is worth noting, however, that there are important limitations in using deep neural networks for insights into neurobiological processes. One of the critical limiting factors of the analyses described here is our reliance on a pre-trained computational model. Deep CNNs have large numbers of parameters, which are typically fit through supervised learning using millions of labeled stimuli. Given the cost of manually labeling this number of stimuli and the far smaller number of stimuli used in a typical neuroscience experiment, it is not feasible to train deep neural networks that are customized for the perceptual processes of interest in every new experiment. Fortunately, neuroscientists can take advantage of the fact that deep neural networks trained for real-world tasks using large, naturalistic stimulus sets appear to learn a set of general-purpose representations that often transfer well to other tasks [25, 50, 51]. Furthermore, the objective functions that these CNNs were trained for all relate to computer-vision goals (e.g., object or scene classification), and, yet, their internal representations exhibit remarkable similarities to those at multiple levels of visual cortex [5-11]. This means that investigators can examine existing pre-trained models for their potential relevance to a cortical sensory process, even if the models were not explicitly trained to implement that process. However, in the analysis of pre-trained models, the architecture, activation function, and other design factors are constrained, and, thus, the results of these analyses cannot be easily compared with alternative algorithmic implementations. An important direction for future work will be the use of multiple models to compare specific architectural and design factors with neural processes, such as the number of model layers, the directions and patterns of connectivity between neurons, the kinds of non-linear operations that the neurons implement, and so on. Nonetheless, we still have much to learn about the information processes of existing CNNs and how they relate to cortical sensory functions, and there is fruitful work to be done in developing techniques that leverage these models for theoretical insights.

Another limitation of this work is that many computer-vision models, and most visual neuroscience experiments, are restricted to simple perceptual tasks using static images. This ignores many important aspects of natural vision that any comprehensive computational model will ultimately need to account for, including attention, motion, temporal dependencies, and the role of memory. Our findings are also limited by the noise ceiling of our neural data. Although the CNN explained as much variance in the OPA as could be expected for any model, there still remains a large portion of variance that can be attributed to noise. This noise arises from multiple factors, including inter-subject variability, variability in cortical responses across stimulus repetitions, the limited resolution of fMRI data, signal contamination from experimental instruments or physiological processes, and irreducible stochasticity in neural activity. Improving the noise ceiling in fMRI studies of high-level visual cortex will be an important goal for future work. Several of these noise-related factors could potentially be mitigated through the use of larger and more naturalistic stimulus sets or through improved data pre-processing procedures. Others, such as inter-subject and inter-trial variability may have identifiable underlying causes that are important for understanding the functional algorithms of high-level visual cortex [52, 53].

In addition to the experimental approaches used here, there are other important avenues of investigation for relating the computations of neural networks to the visual system. For example, CNNs could be used to generate new stimuli that are optimized for testing specific computational hypotheses [17], and then fMRI data could be collected to examine the role of these computations in visual cortex. Another useful approach would be to run *in silico* lesion studies on CNNs to understand the role of specific units within a computational system. Finally, an important direction for future work will be to use the conclusions of experiments on CNNs to build simpler models that embody specific computational principles and allow for detailed investigations of the necessary and sufficient components of information processing in vision.

### Conclusion

An important goal of neuroscience is to understand the computational operations of biological vision. In this work, we utilized recent advances in computer vision to identify an image-computable, quantitative model of navigational-affordance coding in scene-selective visual cortex. By running experiments on this computational model, we characterized the stimulus inputs that drive its internal representations, and we revealed the complex, high-level scene features that its computations give rise to. Together, this work suggests a computational mechanism through which visual cortex might encode the spatial structure of the local navigational environment, and it demonstrates a set of broadly applicable techniques that can be used to relate the internal operations of deep neural networks with the computational processes of the brain.

## METHODS

### Representational similarity analysis

We used RSA to characterize the navigational-affordance information contained in multivoxel fMRI-activation patterns and multiunit CNN-activity patterns.

For the fMRI data, we extracted activation patterns from a set of functionally defined ROIs for each of the 50 images in the stimulus set, using the procedures described in our previous report [19]. Briefly, 16 subjects viewed 50 images of indoor scenes presented for 1.5 s each in 10 scan runs, and performed a category-detection task. Subjects were asked to fixate on a central cross that remained on the screen at all times and press a button if the scene they were viewing was a bathroom. A general linear model was used to extract voxelwise responses to each image in each scan run. ROIs were based on a standard set of functional localizers collected in separate scan runs, and they were defined using an automated procedure with group-based anatomical constraints [19, 54, 55].

For each subject, the responses of each voxel in an ROI were z-scored across images within each run and then averaged across runs. We then applied a second normalization procedure in which the response patterns for each image were z-scored across voxels. Subject-level RDMs were created by calculating the squared Euclidean distances between these normalized response patterns for all pairwise comparisons of images. The squared Euclidean distance metric was used (here and for the other RDMs described below) because several of our analyses involved multiple linear regression for assessing representational similarity. In this framework, the distances from one RDM are modeled as linear combinations of the distances from a set of predictor RDMs. This requires the use of a distance metric that sums linearly [6, 56]. Squared Euclidean distances sum linearly according to the Pythagorean theorem, and when the representational patterns are normalized (i.e., z-scored across units), these distances are linearly proportional to Pearson correlation distances, which we used in our previous analyses of these data [19]. We then constructed group-level neural RDMs for each ROI by taking the mean across all subject-level RDMs. The use of group-level RDMs allowed us to apply the same statistical procedures for assessing all comparisons of RDMs (i.e., fMRI RDMs, navigational-affordance RDM, and CNN RDMs). Furthermore, the use of group-level RDMs, which are averaged across subjects, has the benefit of increasing signal-to-noise and improving model fits for the RSA comparisons.

To construct RDMs for each layer of the CNN, we first ran the experimental stimuli through a pretrained CNN that can be downloaded here: http://places.csail.mit.edu/model/placesCNN_upgraded.tar.gz. We recorded the activations from the final outputs of all linear-nonlinear operations within each layer of the CNN. All layers, with the exception of layer 8, contain thousands of units. We found that the RSA correlations between the layers of the CNN and the ROIs were improved when the dimensionality of the CNN representations was reduced through principal component analysis (PCA; data not shown). This likely reflects the fact that all CNN units were weighted equally in our calculations of representational distances, even though many of the units had low variance across our stimuli, which were all indoor scenes. PCA reduces the number of representational dimensions and focuses on the components of the data that account for the largest variance. We therefore set the dimensionality of the CNN representations to 45 principal components (PCs) for each layer. Our findings were not contingent on the specific number of PCs retained; we observed similar results across the range from 30 to 49 PCs. We z-scored the CNN activations across PCs for each image and calculated squared Euclidean distances for all pairwise comparisons of images.

The neural and CNN RDMs were compared with an RDM constructed from the representations of a navigational-affordance model. To construct this model, we calculated representational patterns that reflected the navigability of each scene along a set of angles radiating from the bottom center of the image (Fig. 1). These navigability data were obtained in a norming study in which an independent group of raters, who did not participate in the fMRI experiment, indicated the paths that they would take to walk through each scene (Fig. 1B) [19]. In our previous report, we combined these navigational data with a set of idealized tuning curves that reduced the dimensionality of the data to a small set of hypothesized encoding channels (i.e., paths to the left, center, and right). Here, however, we used a different approach in which we simply smoothed the navigability data over the 180 degrees of angular bins using an automated and robust smoothing method [57]. This smoothing procedure was implemented using publicly available software from the MATLAB file exchange: https://www.mathworks.com/matlabcentral/fileexchange/25634-fast--n-easy-smoothing?focused=6600598&tab=function. We then z-scored these smoothed data across the angular bins for each image and calculated squared Euclidean distances for all pairwise comparisons of images.

For standard RSA comparisons of two RDMs, we calculated representational similarity using Spearman correlations. The Spearman-correlation procedure assesses whether two models exhibit similar rank orders of representational distances, which allows for the possibility of a nonlinear relationship between the pairwise distances of two RDMs. Nonetheless, we observed similar results using Pearson correlations or linear regressions, and thus our RSA findings were not contingent on the use of a non-parametric statistical test. Bootstrap standard errors of these correlations were calculated over 5000 iterations in which the rows and columns of the RDMs were randomly resampled. This is effectively a resampling of the stimulus labels in the RDM. Resampling was performed without replacement by subsampling 90% of the rows and columns of the RDMs. We did not use resampling with replacement because it would involve elements of the RDM diagonals (i.e., comparisons of stimuli to themselves) that were not used when calculating the RSA correlations [58]. All bootstrap resampling procedures were performed in this manner. Statistical significance was assessed through a permutation test in which the rows and columns of one of the RDMs were randomly permuted and a correlation coefficient was calculated over 5000 iterations. P-values were calculated from this permutation distribution for a one-tailed test using the following formula:

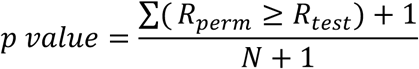

where *R*_*perm*_ refers to the correlation coefficients from the permutation distribution and *R*_*test*_ refers to the correlation coefficient for the original data. All p-values were Bonferroni-corrected for the number of comparisons performed (i.e., the number of ROIs in Fig. 1 and the number of CNN layers in Fig. 2).

We calculated the noise ceiling for RSA correlations in the OPA as the mean correlation of each subject-level OPA RDM to the overall group-level OPA RDM, a measure that reflects the inherent noise in the fMRI data [59]. According to this metric, the best-fitting model for an ROI should explain as much variance as the average subject.

### Commonality analysis

Several analyses involved the use of multiple linear regression and a variance-partitioning procedure to quantify the overlap of explained variance for two predictor RDMs. The multiple linear regression models included two regressors for the predictor RDMs and a third regressor for the constant term. These regressors were used to explain variance in a third RDM, which served as the dependent variable. Thus, the data points for the dependent and independent variables were the pairwise distance measurements of the RDMs. The models were fit using ordinary least squares regression. We quantified the overlap of explained variance for the two predictor RDMs using a procedure known as commonality analysis [31]. This procedure partitions the explained variance of the regression model into the shared and unique components contributed by all regressors. We used this analysis to determine the degree to which the explained variance of one regressor (e.g., the affordance RDM) was shared with a second regressor (e.g., the CNN RDM). We refer to this quantity as shared variance, and we calculated it using the following formula:

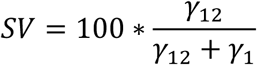

where *SV* is the percentage of the explained variance for regressor X1 that is in common with regressor X2 (see also Fig. 3A). The other variables in this equation refer to components of the overall explained variance 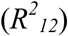:

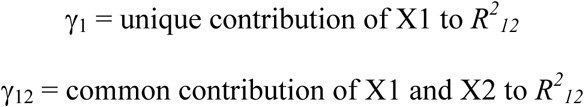

These values are calculated as follows:

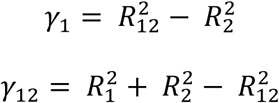

where 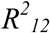 is the explained variance of a regression model with both X1 and X2, 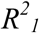 is the explained variance of a model with only X1, and 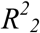 is the explained variance of a model with only X2.

Bootstrap standard errors of this shared-variance metric were calculated over 5000 iterations in which the rows and columns of the RDMs were randomly resampled and the variance-partitioning procedure was applied to the resampled RDMs.

### Analyses of low-level stimulus inputs

We quantified the contribution of specific low-level image features to the RSA effects of the CNN. To do this, we generated new sets of filtered stimuli in which specific visual features or portions of the image were isolated or removed (e.g., color, spatial frequencies, edges at cardinal or oblique orientations, lower or upper portions of the image; Fig. 4-5). These filtered stimuli were passed through the CNN, and new RDMs were created for each layer. We used the commonality-analysis technique described above to quantify the portion of the original explained variance of the CNN that could be accounted for by the filtered stimuli. This procedure was applied to the explained variance of the CNN for predicting both the navigational-affordance RDM and the OPA RDM.

We performed five different stimulus transformations to examine specific classes of image features. The first was a simple transformation of the images from color to grayscale that allowed us to assess the importance of color information. The others reflect two broad categories of low-level image properties: spatial frequencies and contour orientations. To examine the role of spatial frequencies, we created one set of stimuli in which low spatial frequencies were removed from the images (high-pass) and another set in which high spatial frequencies were removed (low-pass). These were created by first converting the images to grayscale, performing a Fourier transform, filtering out a subset of frequencies, and then reconstructing the grayscale images from the filtered Fourier transforms. For the high-pass images, the Fourier spectrum was filtered using a Gaussian filter with a standard deviation set at 0.1 cycles per pixel. A similar approach was used for the low-pass images, with the standard deviation of the Gaussian filter set at 0.0075 cycles per pixel. To examine the role of contour orientations, we created one set of filtered stimuli in which edges at cardinal orientations were emphasized (cardinal) and another set in which edges at oblique orientations were emphasized (oblique). These were created by first converting the images to grayscale and then performing a convolution to extract image contours at cardinal orientations (0 and 90 degrees) or oblique orientations (45 and 135 degrees). The convolution kernels spanned 3 pixels by 3 pixels and are depicted in Fig. S8. Convolutions were performed separately for the two orientations in each set (e.g., 0 and 90 degrees) and a combined output was created by squaring and summing these convolutions and then taking the square root of their sum.

We statistically assessed differences in shared variance across sets of filtered images by calculating confidence intervals on their difference scores. We did this for the following subsets: 1) high-pass minus low-pass and 2) cardinal minus oblique. To do so, we calculated a bootstrap distribution of the difference in shared variance values across each image set over 5000 iterations in which the rows and columns of the RDMs were randomly resampled. From this distribution, we computed the value of the lower 95^th^ percentile for a one-tailed test to determine if the 95% confidence interval was above zero.

We also performed analyses to examine the importance of visual inputs at different positions along the vertical axis of the image. To do this, we generated occluded versions of the stimuli in which everything outside of a small horizontal slice of the image was masked. The exposed slice of the image spanned 41 pixels in height, which was 18% of the overall image height. We used commonality analysis to quantify the portion of the original explained variance of the CNN that could be accounted for by the occluded stimuli. This procedure was repeated with the un-occluded region shifted by a stride of 5 pixels on each iteration until the entire vertical axis of the image was sampled. We generated heat maps of these results by assigning shared variance values to the pixels in each horizontal slice and then averaging the values across overlapping slices.

### Visualizations of high-level feature selectivity

We used a receptive-field mapping procedure in combination with a set of data-driven visualization techniques to gain insights into the complex feature selectivity of units within the CNN. The receptive-field mapping and image-segmentation procedures were based on previously published methods [40]. We mapped the selectivity of individual CNN units across each image by iteratively occluding the inputs to the CNN. First, the original image was passed through the CNN. Then a small portion of the image was occluded with a patch of random pixel values of size 11 pixels by 11 pixels, as in [40]. The occluded image was passed though the CNN, and discrepancies in unit activations relative to the original image were logged. Theses discrepancies were calculated as the absolute value of the difference in activation, which is consistent with the procedure used by Zhou and colleagues [40] (personal communication with Bolei Zhou). On each iteration, the position of the occluding patch was shifted by a stride of 3 pixels. After iteratively applying this procedure across all spatial positions in the image, a two-dimensional discrepancy map was generated for each unit and each image (Fig. 6A). Each discrepancy map indicates the sensitivity of a CNN unit to the visual information across all spatial positions of an image. The spatial distribution of the discrepancy effects reflects the position and extent of a unit’s receptive field, and the magnitude of the discrepancy effects reflects the sensitivity of a unit to the underlying image features.

We generated image segmentations to visualize the scene features that individual CNN units were most sensitive to (Fig. 6B). We first smoothed the discrepancy maps by convolving them with a local averaging filter of 20 pixels by 20 pixels. For each unit, we then selected the 3 stimulus images that generated the largest discrepancy values at any spatial location in the image. We segmented these discrepancy maps by identifying pixels with a discrepancy value equal to at least 10% of the peak discrepancy across all pixels. We generated these visualizations for 50 units in layer 5. These units were chosen based on their unit-wise RSA correlations to the affordance RDM and the OPA RDM (we chose the units with the highest mean correlation to these two RDMs).

We then used t-SNE and k-means clustering to generate a summary visualization and to identify common themes among the scene features that were highlighted by these segmentations [60]. Our goal was to cluster the image segmentations based on the similarity of their high-level scene content. We first created image patches of 81 pixels by 81 pixels centered on the peak discrepancy value for the top 3 images for each unit. We ran these image patches through the CNN and logged the responses in layer 5. These responses were then averaged across the top 3 images for each unit, and t-SNE was used to generate a two-dimensional embedding of all 50 units based on the similarity of their mean response vectors from layer 5. We assigned the units in this embedding to clusters with similar scene features (Fig. 6B and S1-S7). Clusters were identified in a data-driven manner through k-means clustering, with the number of clusters chosen (within the range of 1 to 10 clusters) using the silhouette criterion in the MATLAB function evalclusters.

